# Sustained postsynaptic kainate receptor activation downregulates AMPA receptor surface expression and induces hippocampal LTD

**DOI:** 10.1101/2020.12.05.412981

**Authors:** Jithin D. Nair, Ellen Braksator, Busra P Yucel, Richard Seager, Jack R. Mellor, Zafar I. Bashir, Kevin A. Wilkinson, Jeremy M. Henley

## Abstract

Here we report that sustained activation of GluK2 subunit-containing kainate receptors leads to AMPA receptor endocytosis and a novel form of long-term depression (KAR-LTD_AMPAR_) in hippocampal neurons. The KAR-evoked loss of surface AMPA receptors requires KAR channel activity and is occluded by the blockade of PKC or PKA. Moreover, in acute hippocampal slices, kainate invoked LTD of AMPA EPSCs. These data, together with our previously reported KAR-LTP_AMPAR_, demonstrate that KARs bidirectionally regulate synaptic AMPARs and synaptic plasticity.

## Introduction

Glutamate mediates the overwhelming majority of excitatory neurotransmission and is critical for nearly all aspects of brain function. Ionotropic glutamate receptors comprise NMDA, AMPA, and Kainate (KA) receptor subtypes, whereas there are 8 metabotropic glutamate receptors classified into 3 subfamilies (groups I-III; mGluR1-8) [1]. Each of these receptor subtypes is implicated in forms of long-term potentiation (LTP) and/or long-term depression (LTD) that are fundamental to learning and memory, and their dysregulation is a prominent feature of neurological and neurodegenerative diseases [2].

While the best characterised forms of LTP and LTD are initiated by postsynaptic NMDARs [3, 4], mGluR activation can also lead to both LTP and LTD [5, 6]. Most recently, it has been shown that KARs can induce LTP [7]. Regardless of the induction pathway, changes in postsynaptic AMPARs underpin LTP and LTD expression [8, 9]. Increased surface expression of synaptic AMPARs lead to functional and structural LTP, whereas decreases lead to LTD [10, 11].

KARs can be expressed at both the pre- and postsynaptic sites throughout the brain, where they contribute to the regulation of transmission, neuronal excitability and network activity [12] via coupling to both ionotropic and metabotropic signalling pathways [13–15]. Transient KA stimulation (e.g. 10μM KA for 3 mins in hippocampal neuronal cultures) increases KAR surface expression [16–19] and leads to spine growth [20]. Furthermore, the same stimulation paradigm also enhances AMPAR surface expression, colocalisation with PSD95 and increases AMPAR mEPSCs in acute hippocampal slices via a metabotropic signalling pathway. These data demonstrated a novel, physiologically relevant form of postsynaptic KAR-dependent, NMDAR-independent LTP (KAR-LTP_AMPAR_) [7].

In contrast to transient KA stimulation, sustained KA stimulation (e.g. 10μM KA for 20 mins) causes a long-lasting removal of KARs from the cell surface of hippocampal neurons [19–21]. Here we report that sustained KAR stimulation also evokes a novel form of AMPAR LTD (KAR-LTD_AMPAR_) demonstrating the bidirectional regulation of AMPARs by KAR mediated signalling.

## Results

### Sustained KA treatment reduces surface expression of AMPARs and KARs

Most AMPARs are heterotetramers of GluA1/GluA2 subunits or GluA2/GluA3 subunits [22]. Therefore, we first tested the effects of sustained KA application (10μM KA for 20 mins) on the surface expression of AMPAR subunits GluA1 and GluA2, and the KAR subunit GluK2, in cultured hippocampal neurons. Neurons were pre-treated for 30 minutes with 1μM tetrodotoxin (TTX), to prevent depolarisation-evoked presynaptic glutamate release, and 40μM GYKI 53655, an AMPAR-specific antagonist [23, 24], to prevent direct activation of AMPARs by KA. In parallel, we used a well-established NMDAR-mediated chem-LTD protocol comprising 20μM NMDA and 20μM glycine for 3 minutes, followed by a 17 min incubation in the absence of NMDA, to allow receptor internalisation [25–27]. Surface proteins and whole cell lysates (total protein) were then analysed by Western blotting for GluA2, GluA1, and GluK2 (**Figure 1A**). Surface expression of GluA2, GluA1, and GluK2 were all significantly reduced by both KA and NMDA. There was no change in the surface expression of EGFR, a non-iGluR plasma membrane protein used as a control.

**Figure 1:**
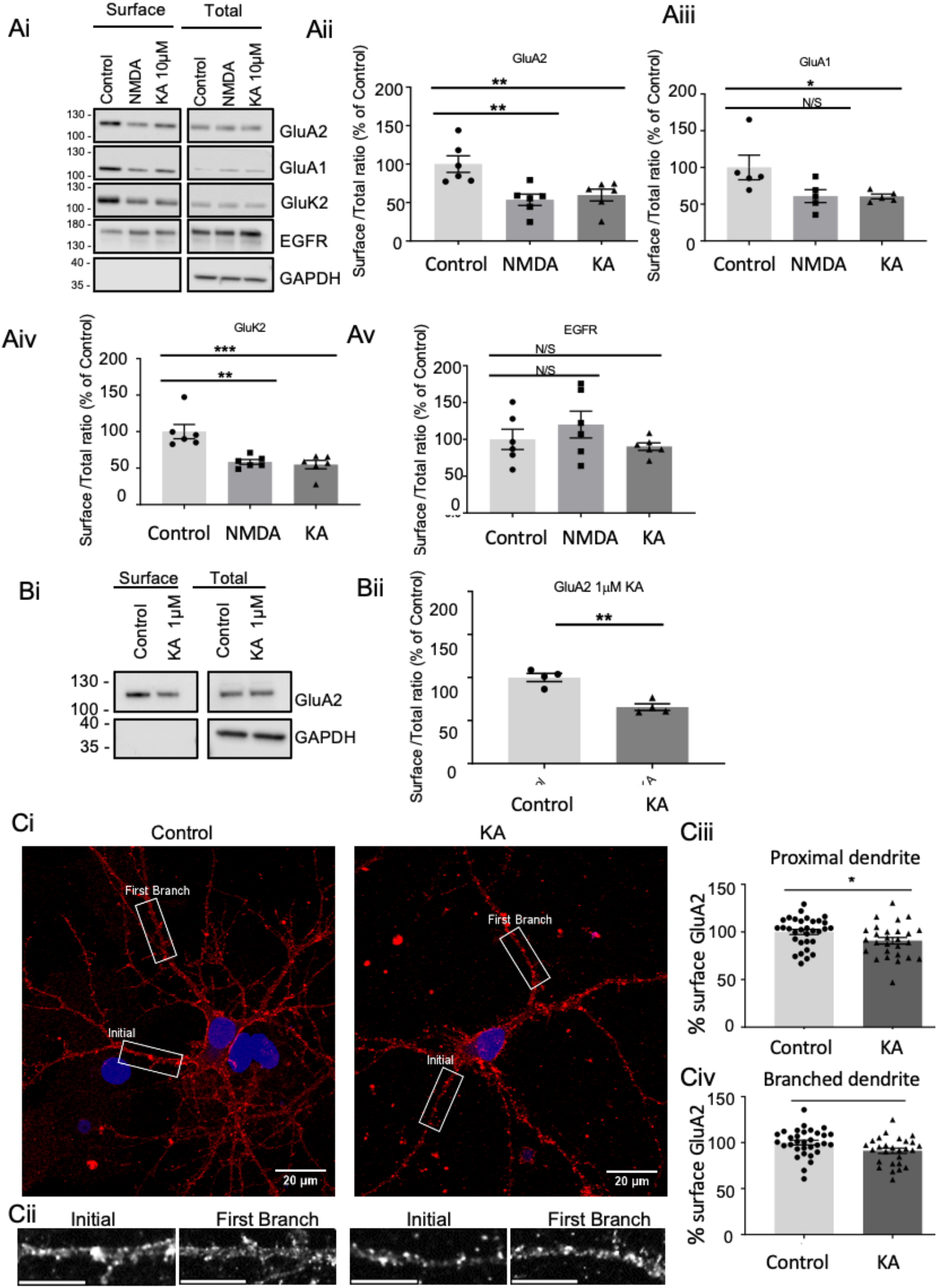
KAR activation reduces surface expression of AMPARs and KARs. DIV 18 cultured hippocampal neurons were pre-treated for 30 minutes with 1μM TTX and 40μM GYKI 53655 before treatment with vehicle or 10μM KA for 20 minutes. For NMDA treatment neurons were pre-treated with 1μM TTX for 30 mins followed by 3 mins of treatment with 20μM NMDA and 20μM glycine. Surface proteins were biotin labelled and isolated by streptavidin pulldown, and lysates and surface fractions Western blotted. **Ai**) Representative Western blots of surface and total levels of GluA2, GluA1, GluK2, and EGFR. EGFR was used as a non-glutamate receptor expressed on the neuronal surface. GAPDH was used as a control to ensure no internal proteins were biotinylated. The surface to total ratio was calculated and expressed as a percentage of the control for (**Aii**) GluA2, (**Aiii**) GluA1, (**Aiv**) GluK2, and (**Av**) EGFR. N=6 experiments from independent dissections, *p<0.05, **p<0.01, ***p<0.001; One-way ANOVA with Dunnett’s multiple comparisons test, error bars = SEM. **B**) 1μM KA effectively reduces surface expression of GluA2-containing AMPARs. (**Bi)** Representative Western blots of surface and total levels of GluA2. (**Bii)** Quantification of the surface to total ratio of GluA2 expressed as a percentage of control. N=4 experiments from independent dissections, **p<0.01, Unpaired t-test with Welch’s correction, error bar = SEM. **C)** Live confocal immunostaining of DIV 18 hippocampal neurons shows a significant reduction in GluA2 surface expression after 10μM KA for 20 minutes. (**Ci**) Representative images of control and KA-treated neurons showing proximal and first branch dendrites. Scale bar = 20μm. (**Cii**) Expanded images of ROIs indicated in boxes in (Ci). Scale bar = 10μm. (**Ciii**) Quantification of the intensity of surface GluA2 staining in proximal dendrites. (**Civ**) Quantification of the intensity of surface GluA2 staining after the first dendritic branch. In all cases n=32 cells, N=3 independent dissections, *p<0.05, Un-paired t-test with Welch’s correction, error bars = SEM.

KA is a partial, weakly desensitising agonist at AMPARs [28]. We therefore tested the effect of 1μM KA, a concentration below the threshold for AMPAR activation [29], in the absence of GYKI 53655, on GluA2 surface expression [30, 31]. As for 10μM KA stimulation, surface levels of GluA2 were significantly reduced by 1μM KA application (**Figure 1B**).

We next performed confocal imaging to monitor GluA2 surface expression in control and KA treated hippocampal neurons. Following pre-incubation with GYKI53655 and TTX, neurons were treated with 10μM KA for 20 mins, followed by live surface labelling with an N-terminal anti-GluA2 antibody. Consistent with the biochemical data, there was a significant reduction in surface GluA2 in both proximal and distal dendrites (**Figure 1C**).

### GluK2-containing KARs mediate decreased AMPAR surface expression

Induction of KAR-LTP_AMPAR_ requires GluK2-containing KARs [7] so we used lentiviral knockdown [32, 33] to determine whether the KAR evoked decreases in surface GluA2 requires the GluK2 KAR subunit. As expected, in neurons infected with control shRNA, KAR activation decreased GluA2 surface expression. However, in GluK2 knockdown neurons there was no KA-induced reduction in surface GluA2 (**Figure 2**).

**Figure 2:**
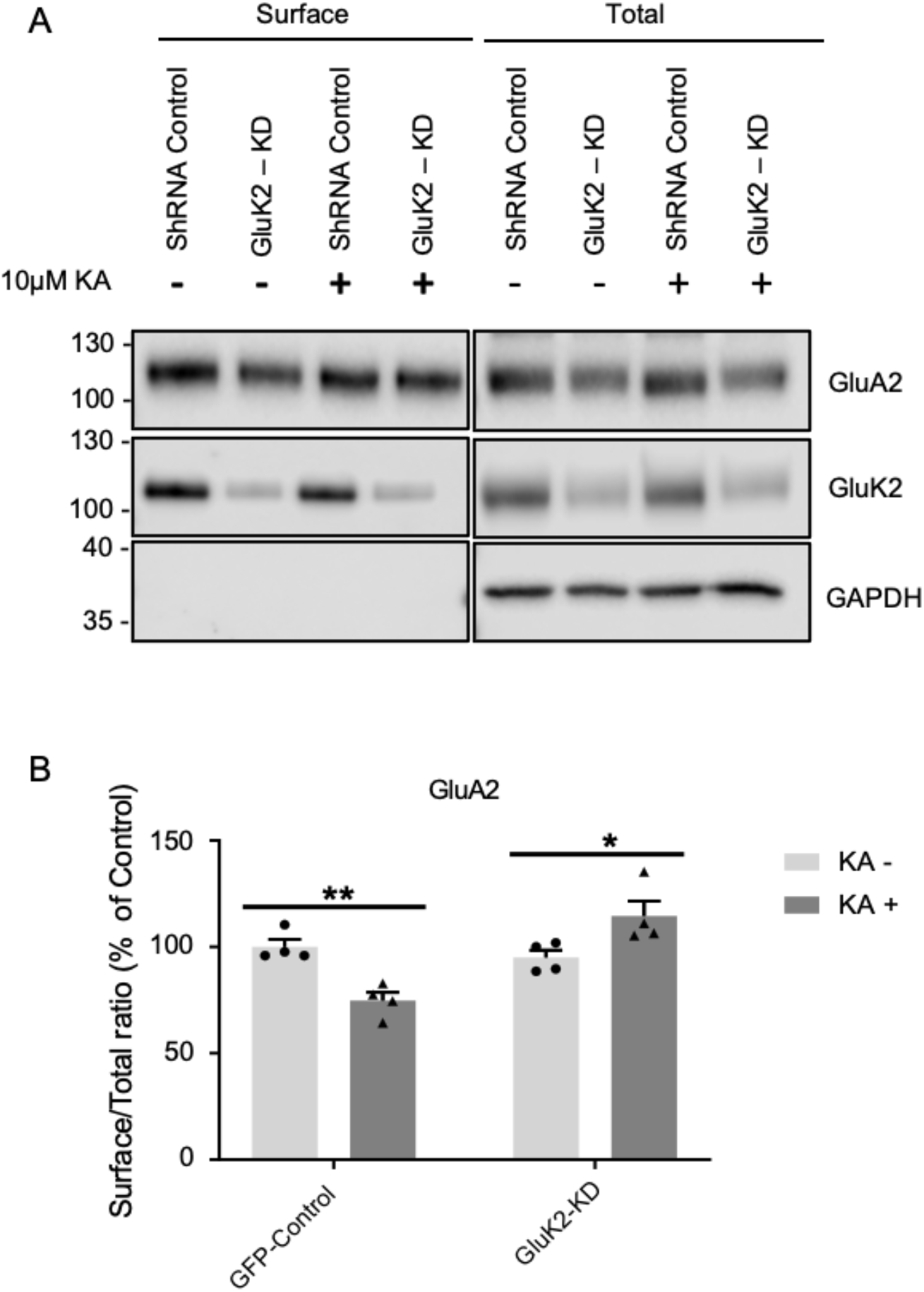
GluK2-containing KARs mediate the reduction in surface expression of AMPARs. **A**) DIV 10 hippocampal neurons were transduced with lentivirus expressing either control or GluK2 shRNA. After 7 days neurons were treated 10μM KA or vehicle and surface proteins isolated and Western blotted. Representative Western blots of surface and total levels of GluA2 and GluK2. **B**) Quantification of the surface to total ratio of GluA2 expressed as a percentage of control. N=4 independent dissections, ns=p>0.05, *p<0.05, **p<0.01, Two-way ANOVA with Dunnett’s multiple comparisons test, error bars = SEM.

### Ionotropic KAR signaling mediates down-regulation of surface AMPARs

To determine the signalling mode that mediates the KAR-induced reduction in surface AMPARs, neurons were pre-treated with 40μM GYKI 53655 and 1 μM TTX for 30 mins, along with the ionotropic KAR blocker UBP 310 (10μM, 30 mins) [7, 34–36] or the G-protein inhibitor pertussis toxin (PTx; 1μg/ml,1hr) [7]. PTx did not prevent the KA-induced reduction in surface GluA2, while UBP 310 did, indicating that KAR channel activity, but not G-protein mediated signalling, is required to downregulate surface AMPARs (**Figure 3A**).

**Figure 3:**
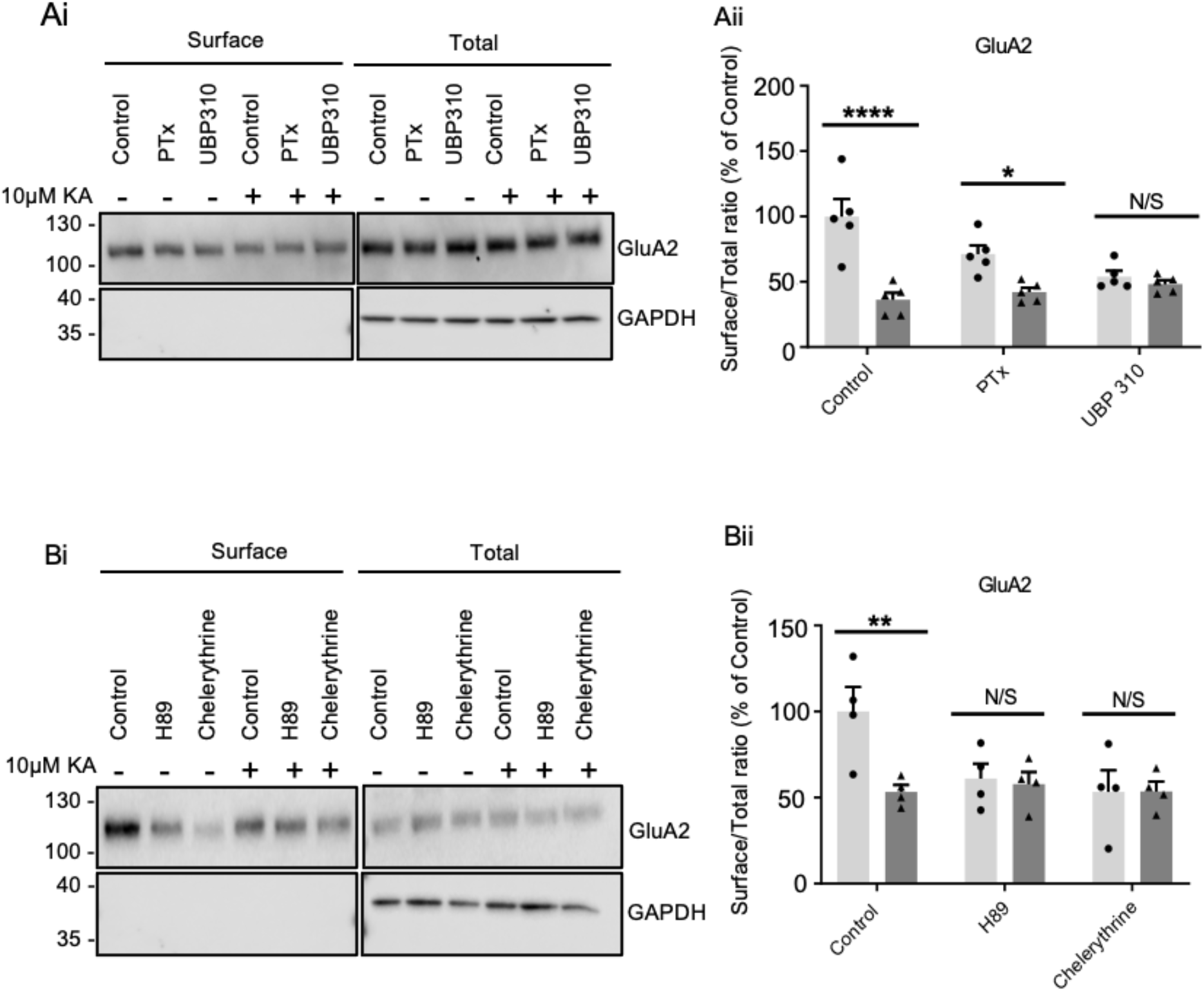
KA-induced decreases in AMPAR surface expression require ionotropic KAR signaling, PKA, and PKC. **A)** DIV 18 hippocampal neurons were pre-treated for 1 h with 1μg/ml PTx, or 30 minutes with 10μM UBP 310, then for 20 min with vehicle or 10μM KA. Surface proteins were biotinylated, isolated and Western blotted. (**Ai**) Representative blot of surface and total levels of GluA2. GAPDH was used as a control to ensure no internal proteins were biotinylated. (**Aii**) Surface to total ratio of GluA2. N=5 independent dissections, *p<0.05, ****p<0.0001, Two-way ANOVA with Dunnett’s multiple comparisons test, error bars = SEM. **B)** As A, except neurons were pre-treated 40μM GYKI 53655 and either 10μM H89 or 5μM Chelerythrine prior to 10μM KA. (**Bi**) Representative blots of GluA2. (**Bii**) Quantification of the surface to total ratio of GluA2 expressed as a percentage of control. N=3 independent dissections, ns p>0.05, *p<0.05, **p<0.01, ***p<0.001; Two-way ANOVA with Dunnett’s multiple comparisons test, error bars = SEM.

### KA regulation of surface GluA2 requires PKA and PKC phosphorylation

We next explored the role of the protein kinases PKA and PKC, which have well-established roles in AMPAR trafficking and KAR signalling [19–21, 37]. Neurons were pre-treated as in Figure 3A but with the PKA and PKC specific inhibitors, H89 (10μM) or chelerythrine (5μM), respectively. Both inhibitors reduced basal surface levels of GluA2 and occluded the KA-induced reduction in GluA2 surface expression (**Figure 3B**).

Together, these data show the KAR-induced decrease in surface GluA2-containing AMPARs requires KAR channel activity, and that PKC or PKA inhibition reduce surface GluA2 and occlude the effect of KAR activation.

### KAR stimulation induces both short-term and long-term synaptic depression

The effects of KA stimulation on AMPAR EPSCs were measured in the CA1 of acute rat hippocampal slices by bath application of KA (1μM, 10 min) in the presence of 50μM picrotoxin and 50μM D-AP5, to block GABA_A_Rs and NMDARs, respectively. AMPAR EPSCs were significantly reduced for 20 mins after KA washout (**Figure 4A**).

**Figure 4:**
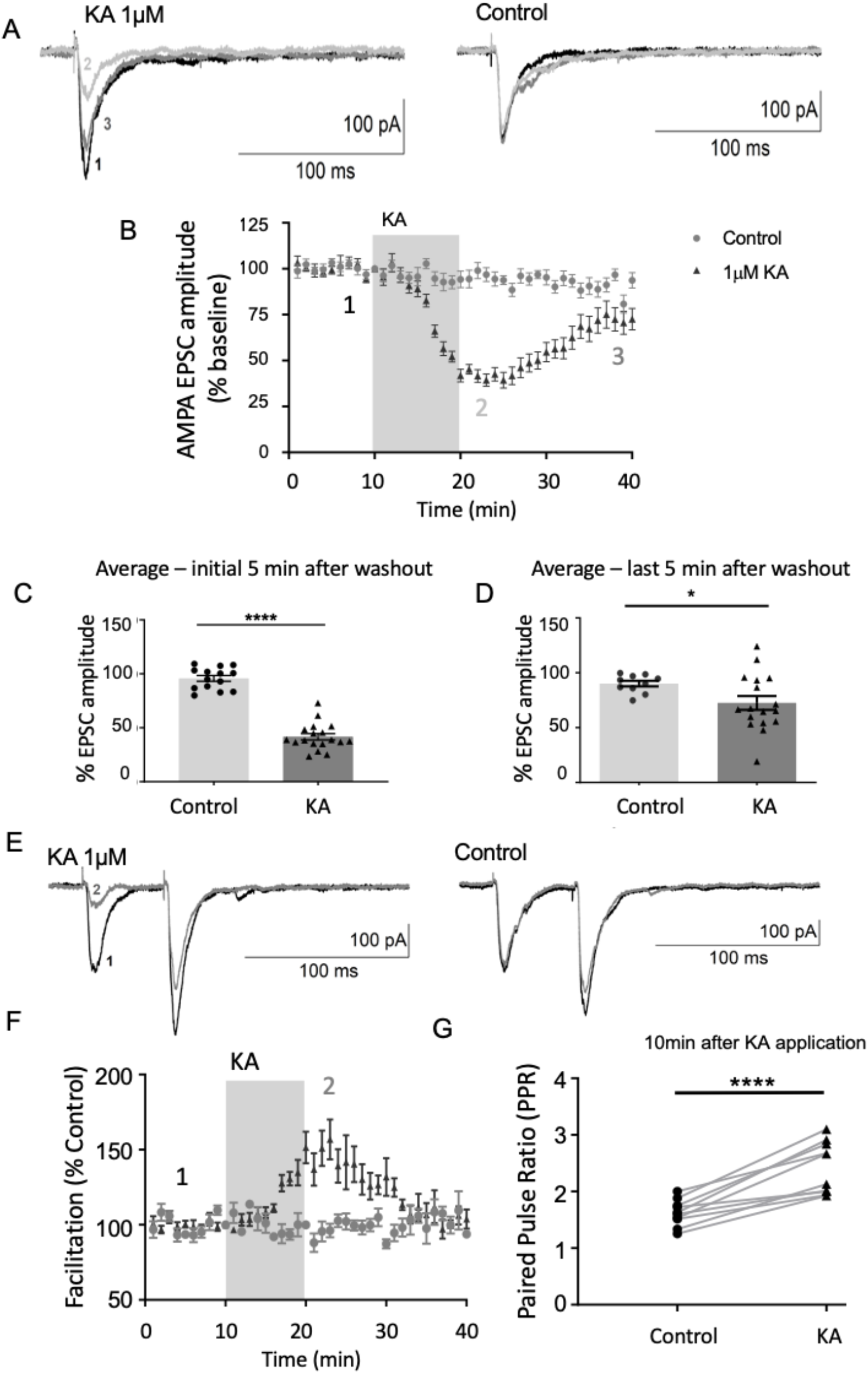
Sustained KAR activation induces depression of AMPAR EPSCs at hippocampal CA1 synapses. **A)** Example AMPAR EPSC traces in the presence or absence of 1μM KA recorded at time points indicated in B in the CA1 region of acute hippocampal slices. **B)** Plot of AMPAR EPSC amplitude. EPSCs were normalised to baseline corresponding to an initial 10 mins prior to KA application. The traces were recorded in the presence of 50μM D-AP5 and 50μM picrotoxin. N=14 and 18 cells for control and KA, respectively, from at least 4 different animals. **C)** Quantification of mean AMPAR EPSC amplitudes in the initial 5 mins after KA washout. Two-way ANOVA with Tukey’s multiple comparisons test, error bars = SEM. **D)** Quantification of mean AMPAR EPSC amplitudes 20 mins after KA washout. N=14 and 18 cells for control and KA, respectively, from at least 4 different animals. Un-paired t-test with Welch’s correction. **E)** Example AMPAR paired pulse EPSC traces in the presence or absence of 1μM KA, recorded at time points indicated in F. **F)** Plot of paired pulse facilitation expressed as a percentage of control after 10 mins of sustained stimulation with 1μM KA. N=10 cells from at least 4 different animals. Error bars = SEM **G)** Quantification of paired pulse ratio after 10 mins of stimulation with 1μM KA. N=10 cells, ns=p>0.05, *p<0.05, **p<0.01,***p<0.001, ****p<0.0001, paired t-test.

We also used paired pulse ratio (PPR) to measure glutamate release [38, 39]. Immediately following KA stimulation PPR was increased, indicating decreased release probability, a characteristic of presynaptic short-term depression (**Figure 4B**) [40]. However, PPR returned to baseline levels within 10-20 mins after KA washout, while AMPAR EPSCs remained depressed. Thus, 1μM KA for 10 min both pre- and postsynaptically-mediate depression of AMPAR EPSCs.

## Discussion

We have reported previously that the surface expression of KARs is subject to bidirectional autoregulation, allowing neurons to adapt their physiological responses to changes in synaptic activation [19]. We further demonstrated that both pharmacological and synaptic activation of GluK2-containing KARs can elicit a novel KAR-dependent, NMDAR-independent, form of hippocampal LTP (KAR-LTP_AMPAR_) through a metabotropic signalling pathway [7]. Here we show that consistent with the bidirectional autoregulation of KARs themselves, GluK2-containing KARs also bidirectionally regulate surface AMPARs. Specifically, prolonged KAR signalling evokes KAR-dependent, NMDAR-independent LTD (KAR-LTD_AMPAR_).

In contrast to KAR-LTP_AMPAR_, which is mediated by metabotropic signalling, KAR-LTD_AMPAR_ requires postsynaptic ionotropic signalling. Interestingly, we also observed a decrease in the basal surface expression of GluA2 in the presence of metabotropic and/or ionotropic inhibitors, suggesting KARs may also be required for maintaining the tonic activity of AMPARs through an as yet unidentified pathway.

Blockade of PKC or PKA reduced GluA2-containing AMPAR surface expression and occluded KAR-evoked LTD, suggesting tonic PKC and PKA activity are required to maintain surface AMPAR levels under control conditions. Moreover, our results raise the possibility that KAR activation potentially causes LTD by reducing PKC/PKA activity.

The physiological/pathological roles of KAR-mediated regulation of AMPAR surface expression at synapses and synaptic plasticity, and how they fit into the larger picture of NMDAR- and mGluR-mediated forms of plasticity remain to be elucidated. However, since KAR abundance and dysfunction is strongly linked to epilepsy [41, 42], it is tempting to speculate that dysfunctional KAR-mediated plasticity of AMPARs could play important roles in neurological diseases.

## Materials and Methods

### Drugs used

| Drugs | Concentration<br>(μM) | Diluted<br>in | Supplier | Catalogue<br>Number |
| --- | --- | --- | --- | --- |
| Chelerythrine | 5 | DMSO | Sigma | C2932 |
| CNQX | 10 | DMSO | Sigma | C127 |
| D-AP5 | 50 | ddH<sub>2</sub>O | Tocris | 0106 |
| Glycine | 20 | ddH<sub>2</sub>O | Sigma | G8790 |
| GYKI 53655 | 40 | ddH<sub>2</sub>O | Tocris | 2555 |
| H89 | 10 | ddH<sub>2</sub>O | Tocris | 2910 |
| Kainate (KA) | 10 | ddH<sub>2</sub>O | Tocris | 0222 |
| NMDA | 20 | ddH<sub>2</sub>O | Tocris | 0114 |
| PTx | 1µg/ml | ddH<sub>2</sub>O | Tocris | 3097 |
| TTX | 1 | ddH<sub>2</sub>O | Tocris | 1069 |
| UBP310 | 10 | DMSO | Tocris | 2079 |
| Picrotoxin | 50 | ddH<sub>2</sub>O | Sigma | P1675 |

### Primary Antibodies used

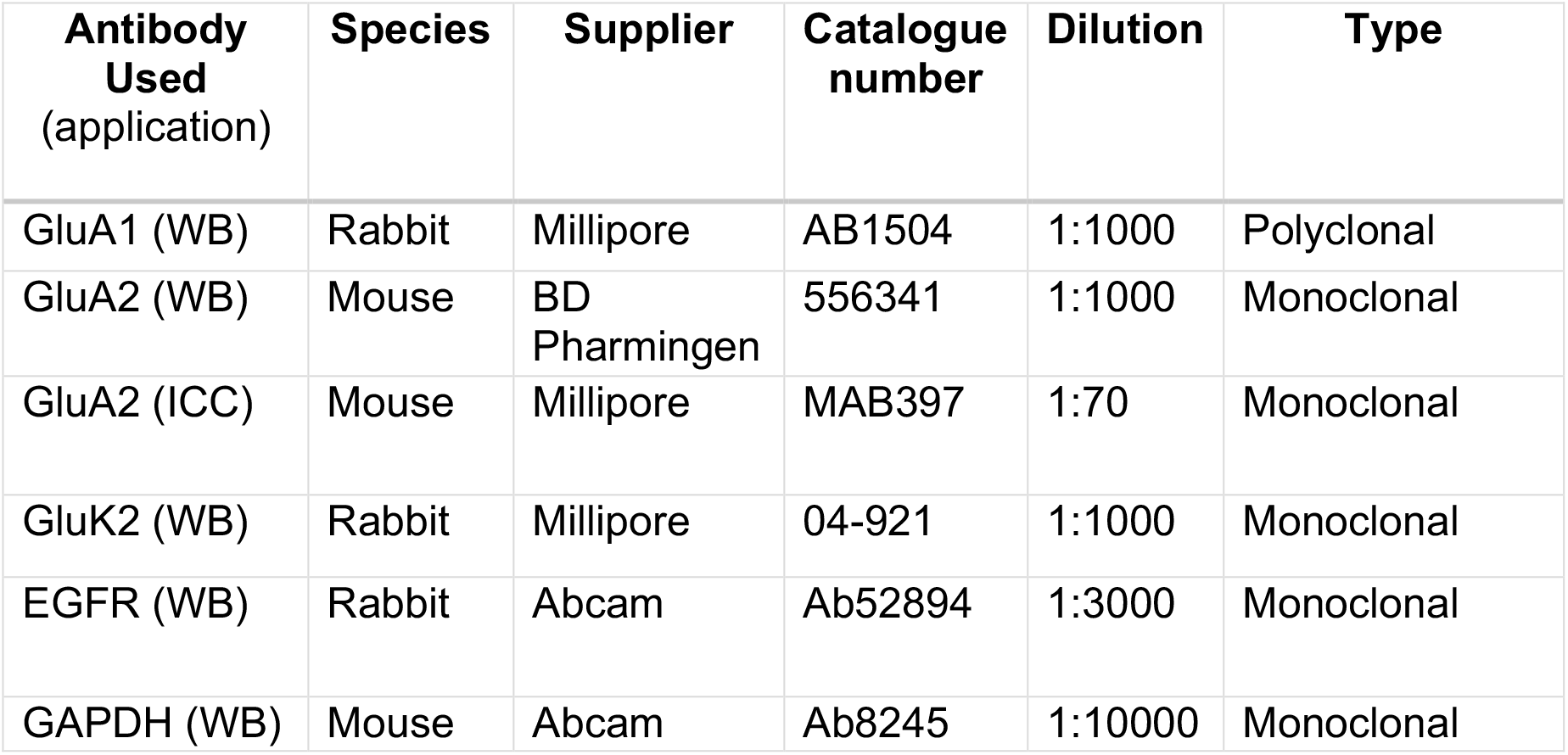

| Antibody Used<br>(application) | Species | Supplier | Catalogue number | Dilution | Type |
| --- | --- | --- | --- | --- | --- |
| GluA1 (WB) | Rabbit | Millipore | AB1504 | 1:1000 | Polyclonal |
| GluA2 (WB) | Mouse | BD Pharmingen | 556341 | 1:1000 | Monoclonal |
| GluA2 (ICC) | Mouse | Millipore | MAB397 | 1:70 | Monoclonal |
| GluK2 (WB) | Rabbit | Millipore | 04-921 | 1:1000 | Monoclonal |
| EGFR (WB) | Rabbit | Abcam | Ab52894 | 1:3000 | Monoclonal |
| GAPDH (WB) | Mouse | Abcam | Ab8245 | 1:10000 | Monoclonal |

### Dissociated primary Neuronal Culture

Primary hippocampal cultures were prepared from E18 Han Wistar rats as previously described [37, 43]. The cells were plated in varying densities for biochemistry and imaging experiments and placed in an incubator for 17-18 days. Neuronal plating media containing Neurobasal (Gibco) supplemented with 10% horse serum, B27, 2mM glutamax, and Penicillin-Streptomycin was changed to feeding media devoid of horse serum after 24 hrs.

### Lentivirus production and transduction

For GluK2 knockdown experiments, shRNA sequences targeting GluK2 were cloned into a modified pXLG3-GFP vector [44] under the control of an H1 promoter. The shRNA target sequences were: Control, non-targeting shRNA: AATTCTCCGAACGTGTCAC; GluK2-targeting shRNA GCCGTTTATGACACTTGGA: The viruses were produced in HEK293T cells as reported previously [44], harvested and added to hippocampal neurons at 9–10 days *in vitro* (DIV) for 7 days before use.

### Sustained KA stimulation

500,000 cells per well of hippocampal neurons were plated in a 6 well culture dish and left in a 37°C incubator until DIV 17-18. On DIV 17, the cells were pre-treated with 1μM TTX (Tocris) and 40μM GYKI 53655 (Tocris), with or without additional drugs as indicated, in Earle’s Buffer (140mM NaCl, 5mM KCl, 1.8mM CaCl_2_, 0.8mM MgCl_2_, 25mM HEPES, 0.9g/L D-Glucose, pH 7.4) and placed back in the incubator for 30 mins. After the pre-incubation, 10μM KA was added to the wells and left for 20 mins at 37°C. Control cells were treated the same way but with vehicle instead of KA.

### Neuronal Surface Biotinylation

Using cell impermeant Sulfo-NHS-SS-Biotin (Thermo Fisher, Cat No.21331), live hippocampal neurons were surface biotinylated post-KA and drug treatment on ice, as described previously [21]. All washes for surface biotinylation were performed with ice cold Earle’s Buffer.

### Immunocytochemistry

Live cell antibody labelling was performed as described previously [45]. Following viral transduction and/or pretreatment with drugs the neurons were stimulated with KA, washed and then incubated with anti-GluA2 N-Terminal Antibody (MAB397, Millipore, 1:70) for 20 mins at 4°C. The cells were washed and fixed in prewarmed 2% formaldehyde (Thermo Fisher) and 2% sucrose for 12 mins. Cells were permeabilized and blocked with 3% BSA containing 0.1% Triton X100 in PBS for 20 mins at RT and labelled with anti-mouse Cy3 antibody (Jackson ImmunoResearch, 1:400) for 1 hr at RT. The coverslips were mounted using Fluoromount-GTM with DAPI (Thermo Fisher) before visualizing on a Leica SP5-II confocal laser scanning microscope.

### Acute slice preparation

Postnatal day 13-15 male and female Han Wistar rats were anaesthetised with 4% isoflurane and decapitated. Brains were rapidly removed and placed in 4°C oxygenated (95% O_2_, 5% CO_2_) sucrose solution (in mM: sucrose 189, D-glucose 10, NaHCO_3_ 26, KCl 3, MgSO_4_ 5, CaCl_2_ 0.1 and NaH_2_PO_4_ 1.25). Parasagittal hippocampal slices 400μm thick were prepared using a vibratome (7000smz-2, Campden Instruments). Slices were kept in a slice holder containing artificial cerebrospinal fluid (aCSF in mM): NaCl 124, NaHCO_3_ 26, KCl 3, NaH_2_PO_4_ 1, MgSO_4_ 1,D-glucose 10 and CaCl_2_ 2; and incubated for 30 mins at 35°C and then for a further 30 mins at room temperature before use.

### Electrophysiology

Hippocampal slices were placed in a submerged holding chamber and perfused with 30°C oxygenated aCSF at 2ml min^-1^. Excitatory postsynaptic currents (EPSCs) of AMPA transmission were evoked at −70mV by stimulating the Schaffer collateral pathway and recorded from CA1 pyramidal neurons. Pyramidal neurons were patch-clamped in the wholecell configuration using borosilicate glass (Harvard Apparatus) electrodes with a resistance of 2-5 MΩ and were backfilled with a solution containing (in mM): CsMeSO_4_ 130, NaCl 8, Mg-ATP 4, Na-GTP 0.3, EGTA 0.5, HEPES 10, QX-314-Cl 5; pH 7.2. The CA3 area of the hippocampal slices was removed using a scalpel blade to minimise epileptic activity. D-AP5 (50μM) and picrotoxin (50μM) were both applied to isolate AMPA-mediated EPSCs. Cells in which the series resistance changed above 20 MΩ or deviated by 20% were discarded.

After a 10 min stable baseline was achieved, 1μM kainic acid was bath applied for 10 mins followed by a 30 min washout period.

Signals were low-pass filtered at 2 kHz and digitised at 10 kHz using an Axopatch 200B amplifier (Molecular Devices) and WinLTP v1.11 acquisition software [46].

### Statistical Analysis

All graphs were generated, and statistical tests performed, using GraphPad Prism version 8.0. Our sample sizes correspond to previously published results and no statistical tests were performed to predetermine the sample size [7, 20]. The details of the statistical tests performed on each experiment are explained in the figure legend along with p-values and error bars. The n (number of cells) and N (number of independent dissections/number of animals) are also mentioned.

## Acknowledgements

We are grateful to the Wolfson Bioimaging Facility (University of Bristol). We also thank the BBSRC (BB/R00787X/1), MRC (MR/L003791/1), Leverhulme Trust (RPG-2019-191), and Wellcome Trust (105384/Z/14/A) for financial support.

## Author contributions

JDN performed all the biochemistry and neuronal live surface staining experiments with help from RS. Confocal imaging was performed by BPY on smaple prepared by JDN. EB performed the electrophysiology guided by ZIB and assistance from JM. DNA constructs were made by KAW. JMH and KAW supervised the study. JDN, KAW, and JMH wrote, and all authors contributed to reading and editing the manuscript.

